# Polioviruses that bind a chimeric Pvr–nectin-2 protein identify capsid residues involved in receptor interaction

**DOI:** 10.1101/162800

**Authors:** Yi Lin, Vincent R. Racaniello

## Abstract

Amino acid changes in the C’C”D region in poliovirus receptor domain 1 disrupt poliovirus binding. To examine further the role of the C’C”D region in poliovirus infection, we substituted this region of Pvr into the corresponding region of a murine homolog, nectin-2. The chimeric receptor, nectin-2^Pvr(c'c"d)^, rendered transformed L cells susceptible to infection with poliovirus P1/Mahoney, but not with polioviruses P2/Lansing and P3/Leon, due to lack of binding. Twenty-four variants of P2/Lansing were selected that replicate in nectin-2^Pvr(c'c"d)^ producing cell lines. Sequence analysis revealed 30 amino acid changes at 28 capsid residues. One change, K1103R, is found in nearly all isolates and is located at one end of the VP1 BC loop. Other alterations are located on the canyon surface, at the protomer interface, and along the perimeter of the canyon south wall. Unlike poliovirus-Pvr binding, the VP1 BC loop is required for infection of cells producing nectin-2^Pvr(c'c"d)^.

## Introduction

The three serotypes of poliovirus initiate infection of primate cells by binding to the poliovirus receptor (Pvr, CD155). Interaction of virus and receptor results in conformational changes in the capsid that ultimately lead to release of the viral RNA genome (Racaniello, 2013). The mechanism by which the receptor interacts with poliovirus has been studied with genetic (Colston and Racaniello, 1994; Colston and Racaniello, 1995; Liao and Racaniello, 1997; Morrison et al., 1994) and structural approaches (Belnap et al., 2000; Bostina et al., 2007; Bubeck et al., 2005; He et al., 2000; He et al., 2003; Strauss et al., 2015; Zhang et al., 2008). These studies describe the virus-receptor interaction and how it leads to release of the viral RNA into the cell.

Poliovirus is a small, nonenveloped RNA virus with an icosahedral shell composed of 60 copies of the capsid proteins VP1, VP2, VP3, and VP4 (Hogle et al., 1985). A prominent peak is centered at each fivefold axis of symmetry and a smaller protrusion is near the threefold axis of symmetry. Clusters of neutralization antigenic sites are located on the highly exposed portions of these surface features (Hogle et al., 1985; Page et al., 1988). A depression or “canyon” that encircles each fivefold axis is the binding site for Pvr, as shown by studies on serotype 1 viruses selected for resistance to soluble receptor (Colston and Racaniello, 1994) or the ability to bind altered receptors (Colston and Racaniello, 1995; Liao and Racaniello, 1997), and solution of structures of the virus-receptor complex (Belnap et al., 2000; Bostina et al., 2007; Bubeck et al., 2005; He et al., 2000; He et al., 2003; Strauss et al., 2015; Zhang et al., 2008).

A member of the immunoglobulin (Ig) superfamily of proteins, the poliovirus receptor (Pvr) is a transmembrane glycoprotein with three extracellular Ig-like domains (Koike et al., 1990; Mendelsohn et al., 1989). The binding site for poliovirus is contained within domain 1 (Koike et al., 1991; Morrison and Racaniello, 1992; Selinka et al., 1991). The results of homolog-scanning mutagenesis studies demonstrate that changes located in the predicted C’-C” and D-E loops, and the C”, D, and G strands significantly disrupt virus binding (Aoki et al., 1994; Bernhardt et al., 1994a; Morrison et al., 1994). Structures of Pvr domain 1 reveal that the region from the C’ strand to D strand (C’C”D region) forms a ridge on one side of the molecule that encompasses the major component of the poliovirus binding site (Belnap et al., 2000; Bostina et al., 2007; Bubeck et al., 2005; He et al., 2000; He et al., 2003; Strauss et al., 2015; Zhang et al., 2008).

To determine if the Pvr C’C”D region is not only necessary but sufficient for poliovirus binding, we constructed a chimeric receptor by substituting this region from Pvr into the corresponding site of a murine homolog of Pvr (Mph/Prr2/Nectin-2), which despite extensive sequence similarity to Pvr does not bind poliovirus (Morrison and Racaniello, 1992). Cells that produce nectin-2^Pvr(c'c"d)^ are susceptible to infection with poliovirus P1/Mahoney, but not with polioviruses of serotypes 2 and 3, which cannot bind to these cells. Binding of poliovirus P1/Mahoney to these cells was weaker than to cells producing Pvr, while serotypes 2 and 3 did not bind at all. Previous studies have also shown that certain amino acid changes in Pvr affect binding of the three poliovirus serotypes differently (Colston, 1995; Harber et al., 1995).

To understand the structural basis for the difference in receptor recognition among the poliovirus serotypes, we isolated 24 variants of poliovirus P2/Lansing that replicate in cells producing the nectin-2^Pvr(c'c"d)^ protein. Amino acid changes that allow P2/Lansing to bind nectin-2^Pvr(c'c"d)^ were identified by sequence analysis and site-directed mutagenesis. One change in capsid protein VP1, K1103R, was found in 21 viral isolates, and its location suggests that the VP1 BC loop regulates interaction of the capsid with the nectin-2^Pvr(c'c"d)^ receptor. Other amino acid changes are located on the canyon floor, at the interface between protomers, and along the outer canyon wall. These alterations increase the binding affinity of P2/Lansing for the chimeric receptor. The locations of these amino acid changes confirm and extend the receptor footprint defined by previous genetic and structural analyses (Belnap et al., 2000; Bostina et al., 2007; Bubeck et al., 2005; Colston and Racaniello, 1994; Colston and Racaniello, 1995; He et al., 2000; He et al., 2003; Liao and Racaniello, 1997; Strauss et al., 2015; Zhang et al., 2008). Furthermore, the results suggest that poliovirus P1/Mahoney contacts the C'C"D region of Pvr, but poliovirus P2/Lansing requires a slightly different contact area. Amino acid changes that allow poliovirus P2/Lansing to bind nectin-2^Pvr(c'c"d)^ receptor might allow contact between the virus and amino acids of nectin-2.

## Materials and Methods

### Cells, viruses, antibodies, and primers

HeLa S3 cells were grown in suspension cultures in Joklik minimal essential medium (Specialty Media) containing 5% bovine calf serum (BCS, Hyclone) and 100 units of penicillin and 100 μg of streptomycin (Pen/Strep, GIBCO) per milliliter. HeLa cells for plaque assays and L tk^-^ cells were plated in Dulbecco’s modified Eagle medium (DMEM, GIBCO) supplemented with 10% BCS, and 100 units-☐g Pen/Strep per milliliter. Stable L cell transformants producing Pvr (20B cells) were additionally maintained with 400 μg of geneticin sulfate (GIBCO) per milliliter. Stable L cell transformants synthesizing nectin-2^Pvr(c’c"d)^ were maintained with 100 mM hypoxanthine, 0.4 mM aminopterin, and 16 mM thymidine (HAT, Sigma).

Working stocks of polioviruses P1/Mahoney, P2/Lansing, and P3/Leon were amplified from seed stocks derived from infectious cDNA clones (Moss et al., 1989; Racaniello and Baltimore, 1981). VP1 BC loop variants of P1/Mahoney, P1/Δ9 and P1/414 virus, were gifts from M. Girard and E. Wimmer (Murray et al., 1988). VP1 BC loop variants of P2/Lansing, R2 1D9 and R2 1D3, were constructed by E. Moss (Moss and Racaniello, 1991).

It should be noted that experiments with poliovirus P2/Lansing were carried out before the 2015 declaration by the WHO that serotype 2 poliovirus has been eradicated; as of mid-2016 the Centers for Disease Control no longer permits work with serotype 2 polioviruses in the US.

Monoclonal antibody 711C reacts specifically with domain 1 of Pvr, while monoclonal antibody 55D reacts with an epitope that bridges domains 1 and 2 of Pvr (Morrison et al., 1994).

DNA primers used for site-directed mutagenesis were purchased from the Columbia DNA Facility and GIBCO. The nucleotide that is changed from the wild type sequence is indicated with underline and bold-face.

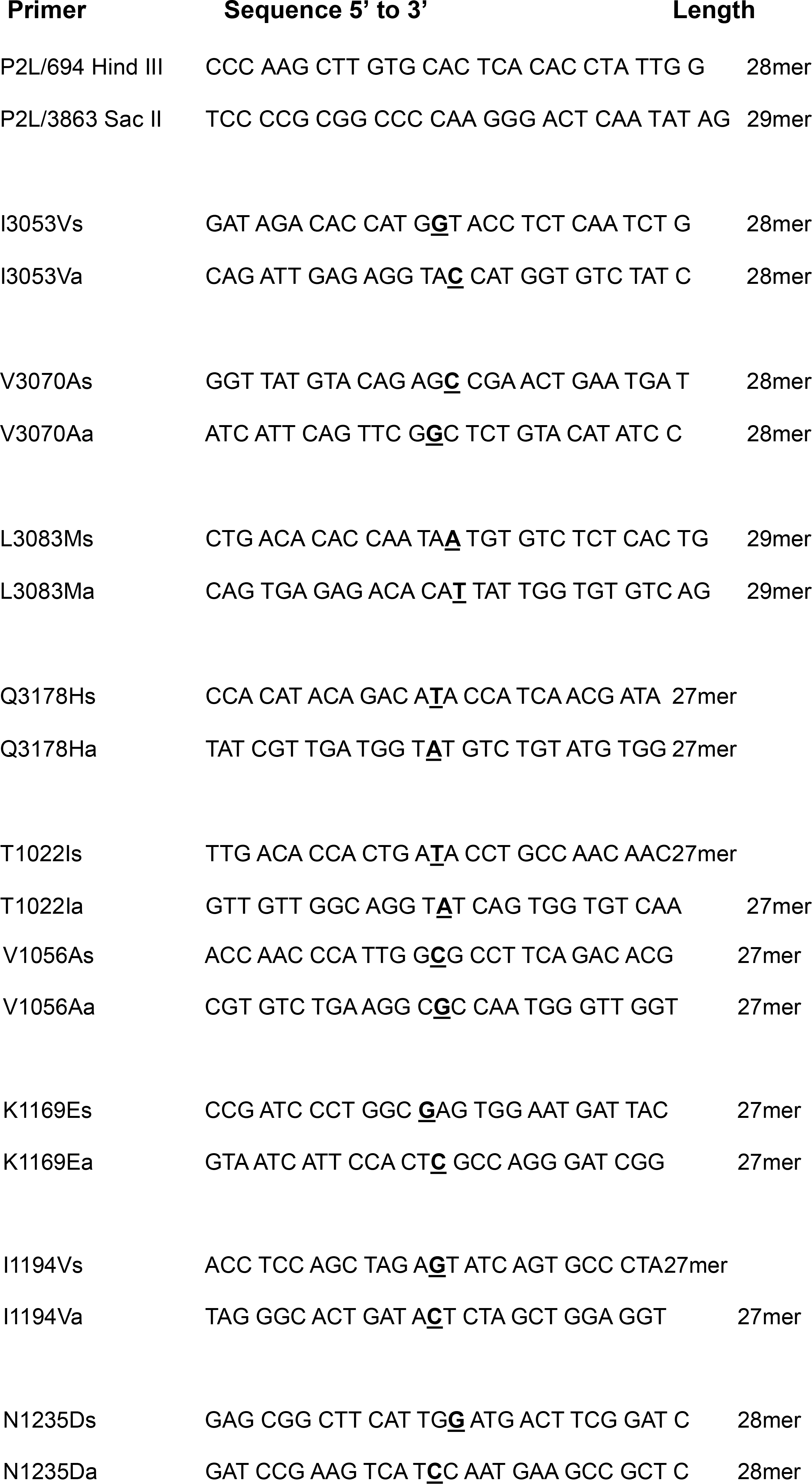

### Construction of nectin-2^Pvr(c'c"d)^ and establishment of stable cell lines

Construction of the nectin-2^Pvr(c'c"d)^ receptor was begun by Mary E. Morrison, using oligonucleotide-directed mutagenesis. The oligonucleotide used for mutagenesis was an 133 nucleotide-long, single-stranded DNA representing the negative-sense sequence of the C’C”D loop-strand region of *pvr* flanked by regions from *nectin-2*:

(3’ TCCACTGGACCGTCGCGGAC **CCACTTAGACCG…CGGTCTGACCCG** CGTCTGGACGCCTACGGTG 5’ The *pvr* sequence is shown in bold). This oligonucleotide was used to prime DNA synthesis on a template consisting of linearized and denatured *nectin-2* DNA in a plasmid. The oligonucleotide was phosphorylated at the 5’-end with T4 polynucleotide kinase (New England Biolabs). The templated was denatured by boiling and slowly cooled in the presence of the oligonucleotide, facilitating its annealing to the template. Negative-strand DNA synthesis was completed with Klenow fragment polymerase (New England Biolabs) and the DNA was recircularized with T4 DNA ligase (New England Biolabs). The plasmid DNA was introduced into *E. coli* DH5αand the desired construct was confirmed by nucleotide sequence analysis. The chimeric *nectin-2* DNA insert was excised via *Xma I* and *Pst I* restriction sites and subcloned into a 5.3 kb partial digestion product of pMEM brSK7+, which contains the *nectin-2* cDNA under the control of an SV40 promoter.

Stable L cell transformants synthesizing Pvr or nectin-2^Pvr(c'c"d)^ were established by the calcium phosphate coprecipitation method as described in (Wigler et al., 1978). Approximately 10μg of plasmid DNA encoding the receptor, and 3 μg of plasmids containing either neomycin resistance gene (pcDNANEO, for Pvr transformants) or the herpesvirus thymidine kinase gene (for nectin-2^Pvr(c'c"d)^ transformants) were coprecipitated with calcium phosphate and added to subconfluent monolayers of L tk^-^ cells in 10 cm plates. The appropriate drug selection was begun after 48 hours incubation. Two weeks post-selection, single colonies were isolated and subcultured in 24 well plates before expanding into 6 cm plates.

Cell surface localization of nectin-2^Pvr(c'c"d)^ could not be detected by available anti-Pvr monoclonal antibodies 711C and 55D (Morrison et al., 1994), probably because the epitope recognized by these antibodies is not present. Cell clones were therefore screened for susceptibility to infection with poliovirus P1/Mahoney. Cultures that produced a tenfold or greater increase in plaque forming units per ml (PFU) of virus at 24 hours post-infection were scored positive for receptor production. Cell clones that produced the greatest increase in viral yield were subjected to a round of clonal sorting by flow cytometry. These clones were rescreened for susceptibility to infection by poliovirus P1/Mahoney, and the highest producing clones were used for this study.

### One-step growth analysis and plaque assays

For analysis of viral replication, subconfluent monolayers of cells plated in 6 cm tissue culture dishes were inoculated with poliovirus at the indicated multiplicity of infection (MOI). After absorption for 30 minutes at 37°C, the plates were washed twice with 4 ml PBS, fresh medium was added and the plates were incubated at 37° C. At the indicated intervals post infection (at t=0 fresh medium is added), cells and supernatant were collected, frozen and thawed three times, and the samples were centrifuged to remove cell debris. Virus titers were determined by plaque assays on HeLa cell monolayers as previously described (Moss et al., 1989).

### Selection of viruses that replicate on cells producing the nectin-2^Pvr(c'c"d)^ receptor

To isolate viruses that replicate on cells synthesizing the nectin-2^Pvr(c'c"d)^ receptor, approximately 10^7^ PFU of P2/Lansing were used to inoculate subconfluent monolayers of L cells synthesizing nectin-2^Pvr(c'c"d)^ (C2 cells) plated in 6 cm tissue culture dishes. Virus was allowed to adsorb for 30 minutes at 37°C before addition of the agar overlay. Approximately three days post-infection, plaques visualized by colormetric “MTT/INT” assay (Shepley et al., 1988) were isolated with 1ml pipets and virus was eluted from agar plugs in dilution buffer (PBS, 0.2% BCS) overnight at 4°C. Two additional rounds of plaque purification on C2 cells were performed for each isolated virus before seed stocks were established.

### Cloning and sequencing of the genomic region encoding the viral capsid

Poliovirus RNA was isolated from virus isolated by ultracentrifugation, and reverse-transcribed according to methods previously described (La Monica et al., 1987). Amplification of the viral capsid coding region by PCR was accomplished with Vent DNA polymerase (NEB). Amplified products from PCR were phosphorylated with T4 polynucleotide kinase (NEB) and cloned via the *EcoR V* restriction site into pBlueScript (Stratagene). For a majority of viruses, cloning was done in duplicates for each isolate. Nucleotide sequence of the capsid coding DNA was determined by the dideoxy method with T7 Sequenase 2 DNA sequencing kit (Amersham Pharmacia) and AmpliCycle sequence kit (Perkin Elmer).

### Site-directed mutagenesis

Several methods were employed to introduce single or double mutations into the genome of poliovirus P2/Lansing. In some cases it was possible to isolate cloned DNA fragments containing only one or two mutations, and substitute the DNA into the cloned DNA of P2/Lansing. For most mutations, either the recombinant PCR approach (Colston and Racaniello, 1995) or the QuickChange Site-Directed Mutagenesis system was used (Stratagene).

### In vitro RNA synthesis and transfection

RNA was transcribed from cloned DNA templates with T7 RNA polymerase (Moss et al., 1989). *In vitro* synthesis of RNA was performed at 37°C for 30 minutes in 50μl of transcription reaction (2 μg of cDNA plasmid, 1 mM each NTP, 50 U RNAsin (Promega), 0.5 μg/ml of BSA (RNAse and DNAse free; Boehringer Mannheim), 5 mM dithiothreitol, 40 mM Tris-Cl (pH 8), 15 mM MgCl_2_, and 35 U of T7 RNA polymerase (Pharmacia). The reaction mixtures were used to transfect HeLa cell monolayers in 10 cm plates with DEAE-dextran as a facilitator (La Monica et al., 1987; McCutchan and Pagano, 1968).

### Metabolic labeling of virus proteins with ^35^S-methionine

Poliovirus was isotopically labeled according to methods previously described (Morrison et al., 1994). Three 15 cm plates of semi-confluent HeLa cells were infected at high multiplicity of infection. After a 30 minute adsorption period, growth medium was added, and infection was allowed to proceed for 90 minutes at 37°C. The infected monolayers were washed twice with 10 ml PBS, and replaced with methionine-free DMEM supplemented with dialyzed BCS and 25 μCi/ml ^35^S-methionine (New England Nuclear). After an overnight incubation, cell lysates were collected, clarified by centrifugation at 40,000 rpm at 10°C for 90 minutes in an SW41 rotor. The virus pellet was overlaid with 200 μl NTE buffer (10mM NaCl, 50mM Tris-HCl [pH 8.0], 10 mM EDTA) with 0.5% SDS. and incubated overnight at room temperature. The resuspended virus was centrifuged through a 7.5 to 45% sucrose gradient in NTE at 40,000 rpm at 10°C for 75 minutes in a SW41 rotor. Gradients were fractionated from the bottom with a glass capillary tube, and the radioactivity of each fraction was determined by liquid scintillation counting. Peak fractions were pooled and dialyzed against PBS. The number of viral particles per ml was calculated from the absorbance at 260 nm, multiplied by the correction factor 9.4 x 10^12^ particles per ml (Rueckert, 1996). Purified virus was adjusted to 5 mg/ml bovine serum albumin (BSA, fraction V, Sigma) and stored in 150 to 250 μl aliquots at –70°C.

### Poliovirus binding assays

Virus binding assays were performed with cell monolayers on 6-well plates. Plates were prepared by seeding cells detached with enzyme-free cell dissociation buffer (GIBCO) into duplicate wells at varying density a day before the experiment. After the removal of the medium, cell monolayers were washed twice with PBS, overlaid with labeled virus (2.7 x 10^10^ particles) in binding buffer (DMEM, 20% Hepes (GIBCO), 10% BCS), and incubated at the determined temperature. After the time indicated in the figure, unattached viral particles were removed, the cell monolayer were washed and lysed with 0.1N NaOH, and the radioactivity was quantitated by liquid scintillation counting. One plate was set aside for determination of the cell number in the wells.

## Results

### Isolation of L cell lines producing Pvr-nectin-2 chimera

Previous studies have demonstrated that mutations located within the predicted C’, C”, and D strands and intervening loops of Pvr domain 1 disrupt poliovirus binding; this region is a principal contact site in the virus-receptor complex (Aoki et al., 1994; Belnap et al., 2000; Bernhardt et al., 1994b; Bostina et al., 2007; Bubeck et al., 2005; He et al., 2000; He et al., 2003; Morrison et al., 1994; Strauss et al., 2015; Zhang et al., 2008). To determine whether this sequence is sufficient to confer poliovirus binding to another Ig-like molecule, it was substituted in the corresponding region of nectin-2. The resulting protein, called nectin-2^Pvr(c'c"d)^, consists of the nectin-2 molecule in which amino acids 30-70 have been substituted with the sequence from Pvr (Figure 1). Stable mouse L cell lines that synthesize nectin-2^Pvr(c'c"d)^ were established. Two cloned cell lines, C1 and C2, were selected for this study. Mouse L cells do not synthesize Pvr (Mendelsohn et al., 1989).

**Figure 1:**
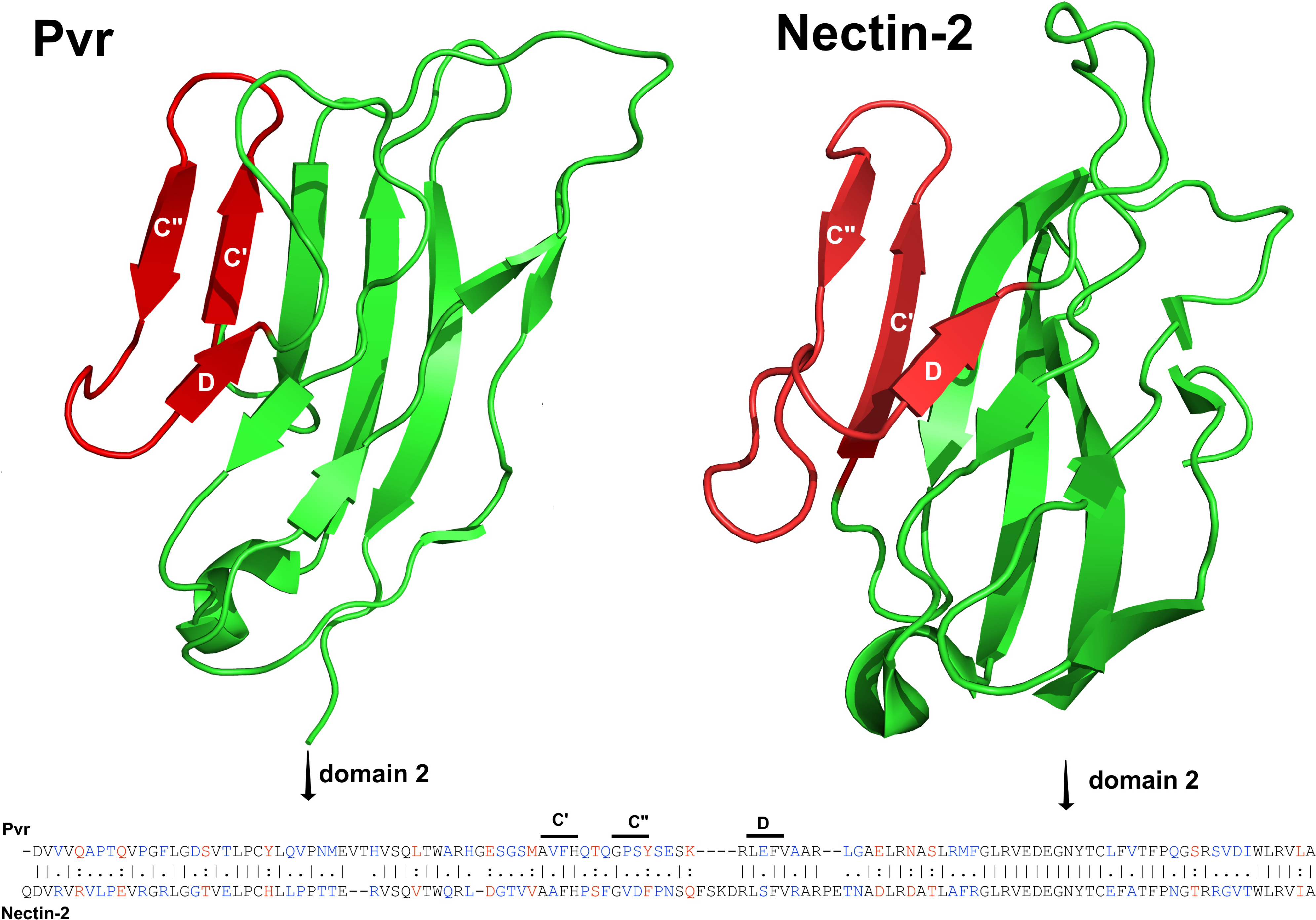
Domain 1 of Pvr and nectin-2. Left, domain 1 of Pvr, with the C’, C”, and D β-strands colored red. These sequences (amino acids 69-100) replaced nectin-2 sequences in in nectin-2^Pvr (c’c”d)^. Model based on pdb file 3j9f (Strauss et al., 2015). Right, domain 1 of nectin-2, with the C’, C”, and D β-strands (amino acids 69-100) that were substituted with sequences from Pvr in nectin-2^Pvr (c’c”d)^ colored red. Based on pdb file 4fmk (Harrison et al., 2012). Bottom, amino acid alignments of domain 1 of Pvr and nectin-2. Identical amino acids are shown in black with a vertical bar; conservative differences are in red with a colon, and semi-conservative differences are in blue with a period. The C’, C”, and D β-strands are labeled. Sequences were aligned using EMBOSS needle (Rice et al., 2000).

### P1/Mahoney replicates in L cells producing nectin-2^Pvr(c'c"d)^

Single-step growth analyses were conducted with representatives of each of the three poliovirus serotypes on L cells synthesizing Pvr (20B cells), nectin-2^Pvr(c'c"d)^ (C1, C2 cells), or recombinant nectin-2 molecules in which domain 1 (E2 cells) or domains 1+2 (H52 cells) are substituted with the corresponding domains from Pvr (Morrison et al., 1994). As expected, all three serotypes of poliovirus replicated in 20B, E2 and H52 cells (Figure 2). Poliovirus P1/Mahoney replicated in C1 and C2 cells, but the kinetics of virus production were different from that in 20B cells (Figure 2a). In 20B cells, after an initial eclipse phase, virus titers increased exponentially to a plateau at approximately 12 hours post-infection. In C1 and C2 cells, the increase in viral titers was more gradual and linear, and did not reach a plateau. Nonetheless, the final titers in C1, C2 and 20B cells were similar. Cytopathic effects typical of poliovirus-infected cells developed in C1 and C2 cells infected with poliovirus P1/Mahoney, although at a slower rate than in 20B cells (unpublished data).

**Figure 2:**
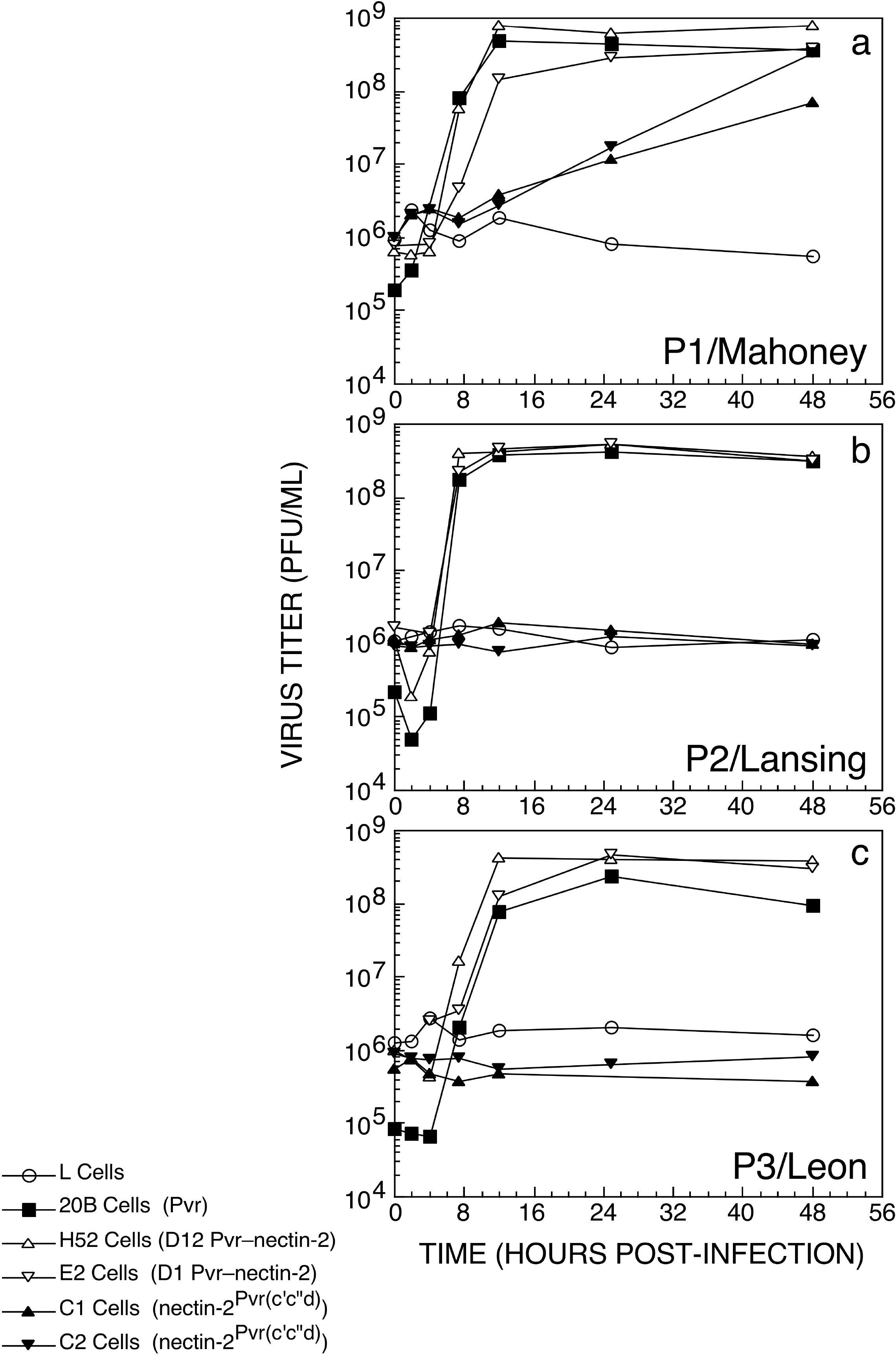
Growth curve of polioviruses in cells producing nectin-2^Pvr (c’c”d)^. ⃝ L cell, ■ 20B cell (Pvr), ΔH52 cell (D12 Pvr-nectin-2), 14∇ E2 cell (D1 Pvr-nectin-2), ▲ C1 and ▼ C2 cell (nectin-2^Pvr (c’c”d)^ monolayers were infected with each of the three serotypes of poliovirus at an MOI of 20, P1/Mahoney (a), P2/Lansing (b) and P3/Leon (c). Total virus titer at the indicated times post-infection was determined by plaque assay on HeLa cells.

In contrast to these results, C1 and C2 cells were not susceptible to infection with poliovirus P2/Lansing or P3/Leon (Figure 2). Virus titers observed at different times after infection of C1 and C2 cells with these viruses resembled those observed after infection of nonsusceptible L cells. These results suggest that in contrast to P1/Mahoney, P2/Lansing and P3/Leon cannot recognize the nectin-2^Pvr(c'c"d)^ receptor.

### P1/Mahoney binds to chimeric Pvr-nectin-2 cells

To determine if infection of C2 cells correlates with attachment, binding assays were performed. All three serotypes of poliovirus bound to 20B cells, but not to L cells (Figure 3). At the highest concentration of cells used, approximately 20% of the input P1/Mahoney virus bound to C1 or C2 cells, whereas 60% was bound by 20B cells (Figure 3). In contrast, little or no binding of P2/Lansing or P3/Leon to C1 or C2 cells was observed. We conclude that P1/Mahoney can bind to the nectin-2^Pvr(c'c"d)^ receptor while P2/Lansing and P3/Leon cannot.

**Figure 3:**
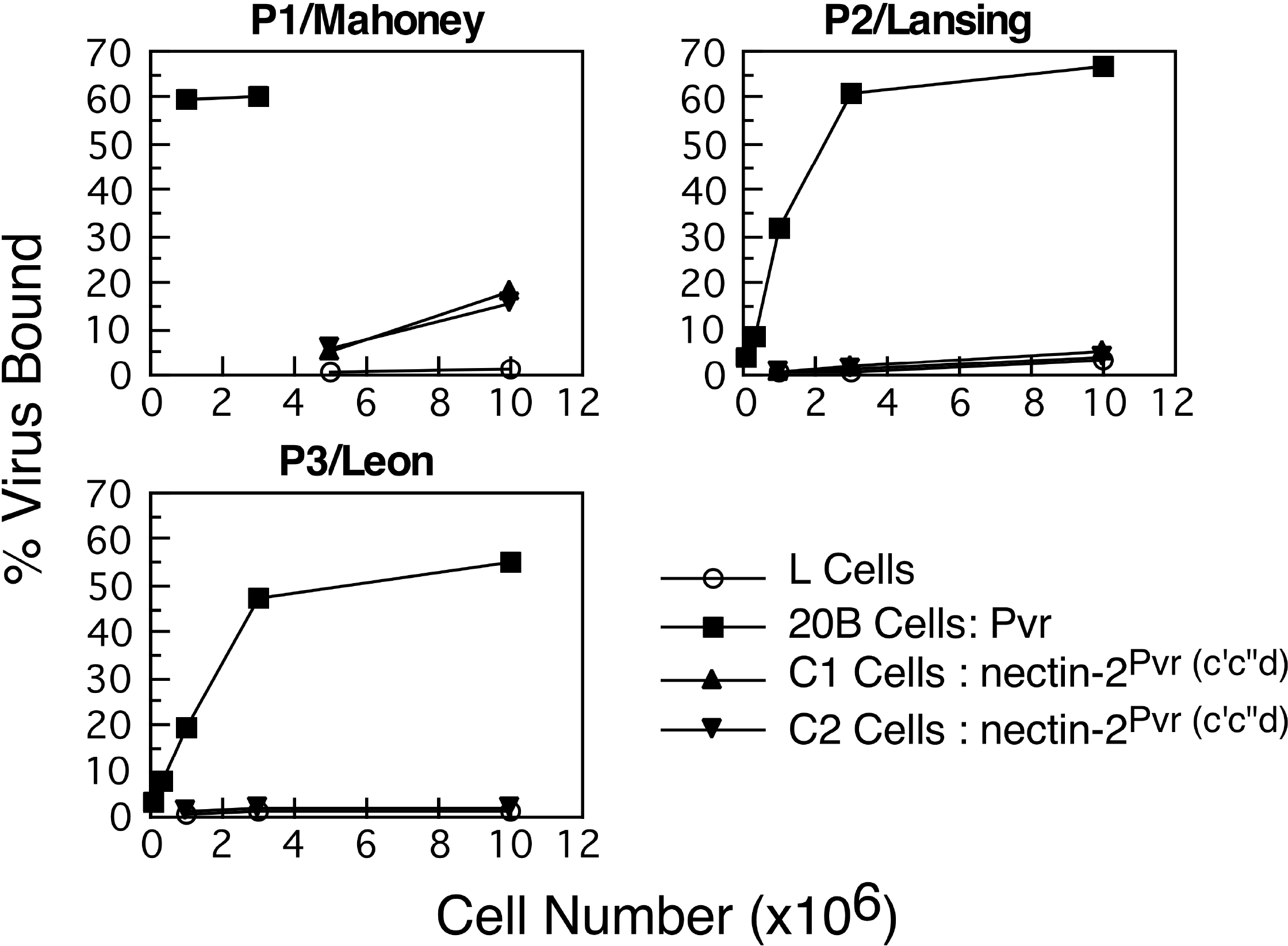
Binding assay of poliovirus on cells producing Pvr or nectin-2^Pvr(c’c”d)^ Radiolabeled poliovirus was incubated with ⃝ L cells, ■ 20B cells, and ▲ C1 and ▼C2 cells overnight at 4°C. Specific binding is expressed as a percentage of the total input virus.

### Selection of P2/Lansing mutants that replicate in cells synthesizing nectin-2^Pvr(c'c"d).^

Poliovirus P1/Mahoney formed plaques on monolayers of C2 cells, although the plating efficiency was about 100 to 1000 fold lower than on 20B cells (Table 1). Virus stocks with a titer of 10^9^ PFU/ml on 20B cells produced an average titer of 10^6^ PFU/ml on C2 cells. An additional day of incubation was necessary for plaques to develop on C2 cells. The reduced efficiency of plating and the longer incubation period required for plaque formation are consistent with defects in the viral life cycle observed in the replication assays (Figure 2a).

**Table 1.** Virus titer from cells producing Pvr or nectin-2^Pvr(c’c”d)^.

| Virus Titer (PFU/ML)<sup>a</sup> |  |  |
| --- | --- | --- |
| Virus | 20B Cells<sup>b</sup> | C2 Cells<sup>c</sup> |
| P1/Mahoney | 3.2 x 10<sup>9</sup> | 6.3 x 10<sup>6</sup> |
| P2/Lansing | 1.0 x 10<sup>9</sup> | 4.0 x 10<sup>2</sup> |
| P2/2B1.1<sup>d</sup> | 2.5 x 10<sup>8</sup> | 5.0 x 10<sup>6</sup> |
| P2/3B1.1 | 3.2 x 10<sup>8</sup> | 6.3 x 10<sup>6</sup> |
| P2/4A1.1 | 1.3 x 10<sup>9</sup> | 6.3 x 10<sup>6</sup> |
| P2/5B1.1 | 1.0 x 10<sup>9</sup> | 5.0 x 10<sup>6</sup> |
| P2/9B1.1 | 1.0 x 10<sup>8</sup> | 2.0 x 10<sup>7</sup> |
| P2/10A1.2 | 2.0 x 10<sup>8</sup> | 2.0 x 10<sup>7</sup> |
| P2/12D1.3 | 1.0 x 10<sup>9</sup> | 1.6 x 10<sup>7</sup> |

<sup>a</sup> determined by plaque assays.
<sup>b</sup> mouse L cells synthesizing Pvr.
<sup>c</sup> mouse L cells synthesizing nectin-2<sup>Pvr(c'c''d)</sup>.
<sup>d</sup> independent viruses isolated by plaque assays of P2/Lansing on C2 cells.

The efficiency of plaquing of poliovirus P2/Lansing in C2 cells was extremely low (Table 1). Viral stocks with a titer of 10^9^ PFU/ml on 20B cells yielded 10^1^ to 10^3^ PFU/ml on C2 cells. To determine whether these plaques were produced by mutant viruses that can replicate in C2 cells, virus was eluted from isolated plaques, subjected to two additional rounds of selection and plaque purification on C2 cells, and viral stocks were generated and titrated on 20B and C2 cells. The titers of the virus isolates on C2 cells were about 10^4^ to 10^5^ fold greater than the titer of the parental virus (Table 1), indicating that they are mutants that can replicate in C2 cells. The titers of the mutant viruses on 20B cells were similar and resembled that of the parent virus. Because the mutant viruses can infect both 20B and C2 cells, they have an expanded host range.

### Identification of capsid alterations in mutants of P2/Lansing that replicate in cells producing nectin-2^Pvr(c'c"d)^

We hypothesized that the mutants of P2/Lansing selected on C2 cells contain amino acid changes in the viral capsid that allow them to recognize the nectin-2^Pvr(c'c"d)^ receptor. To identify the responsible mutations, DNA encoding the capsid coding sequence was molecularly cloned and its nucleotide sequence was determined. Thirty different amino acid changes at 28 different positions were identified from a total of 24 independent virus isolates (Table 2). Many viruses contained one to three mutations, although viruses with four to seven changes were also identified. Six viruses were identified with only one amino acid substitution in the capsid (K1103R, S1239T, or N3059Y).

**Table 2.**
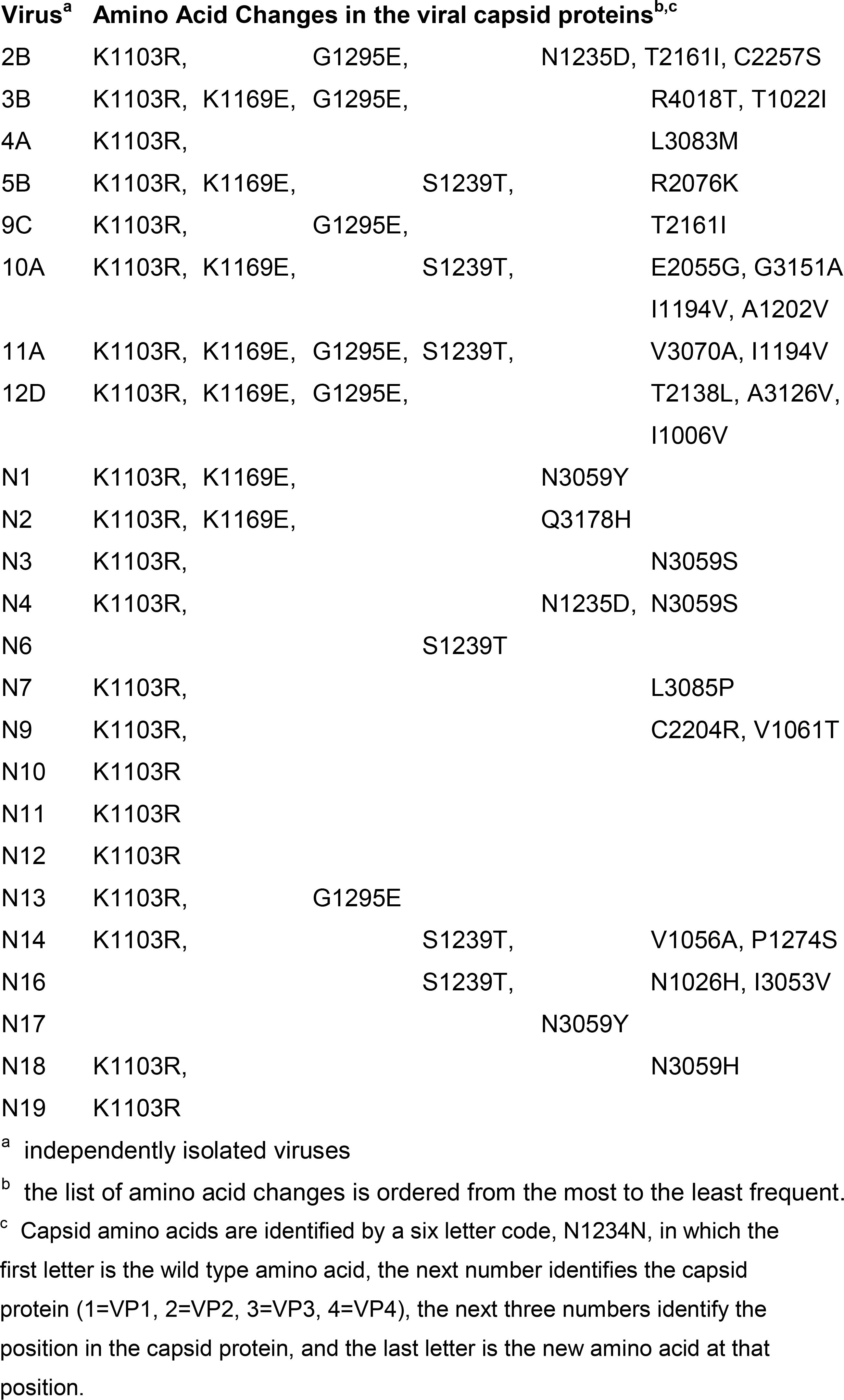
Summary of amino acid changes in P2/Lansing mutants selected on cells synthesizing nectin-2^PVR(c’c”d)^

Some mutations appeared in multiple virus isolates (Table 3). The most common change, K1103R, was identified in 21 viruses. Other frequently occurring substitutions were K1169E, S1239T, and G1295E, each identified in six to seven isolates. Three different amino acid changes (N to H, S or Y) were found at a single position, residue 59 in VP3. Although the frequently occurring changes were often on the same viral capsid, there was no distinct pattern of co-localization (Table 2).

**Table 3.**
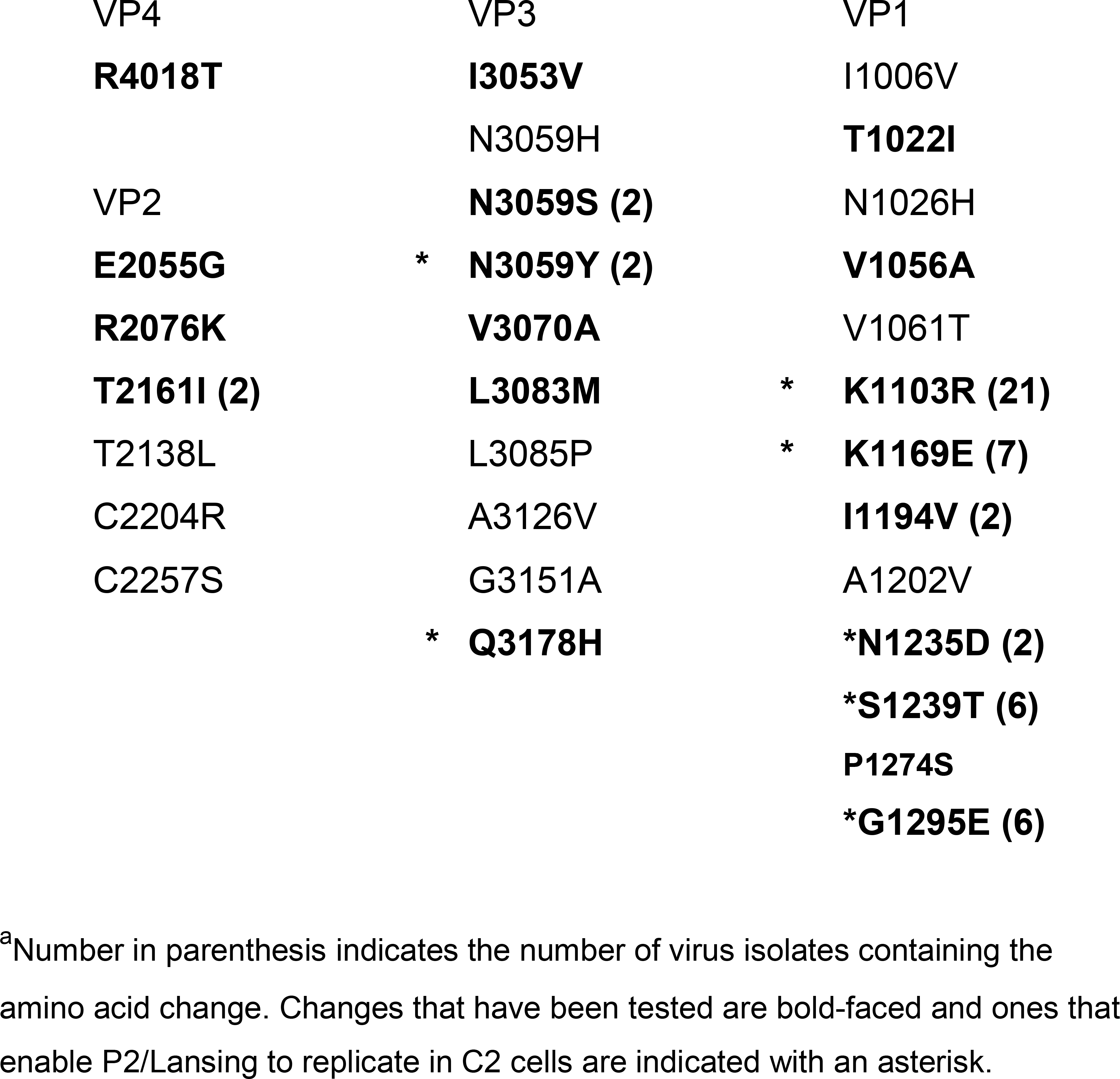
Frequency and distribution of amino acid changes in P2/Lansing mutants selected on cells synthesizing nectin-2^Pvr(c’c”d)^.^a^.

### Identification of seven amino acid changes that enable P2/Lansing to replicate in cells synthesizing nectin-2^Pvr(c'c"d)^

Site-directed mutagenesis was used to determine the effect of single and double amino acid substitutions on the ability of P2/Lansing to replicate in cells producing nectin-2^Pvr(c'c"d)^. Eighteen different substitutions at 17 residues, either individually or in combination with the mutation K1103R, were introduced into the viral genome. Not all changes identified in the original viruses were tested; we selected those which occurred most frequently, or which are located in areas of the viral capsid believed to be important for receptor recognition (Strauss et al., 2015). The titers of the resulting viruses were determined on 20B and C2 cells.

All viruses with single or double amino acid changes plaqued efficiently in 20B cells and produced titers within one log_10_ PFU/ml of P2/Lansing (Figure 4). Single substitutions that enable the virus to replicate in C2 cells were K1103R, K1169E, N1235D, S1239T, G1295E, N3059Y, and Q3178H. Viruses with the changes K1103R, N1235D, S1239T, N3059Y, or Q3178H yielded higher titers on C2 cells than viruses with K1169E or G1295E.

**Figure 4:**
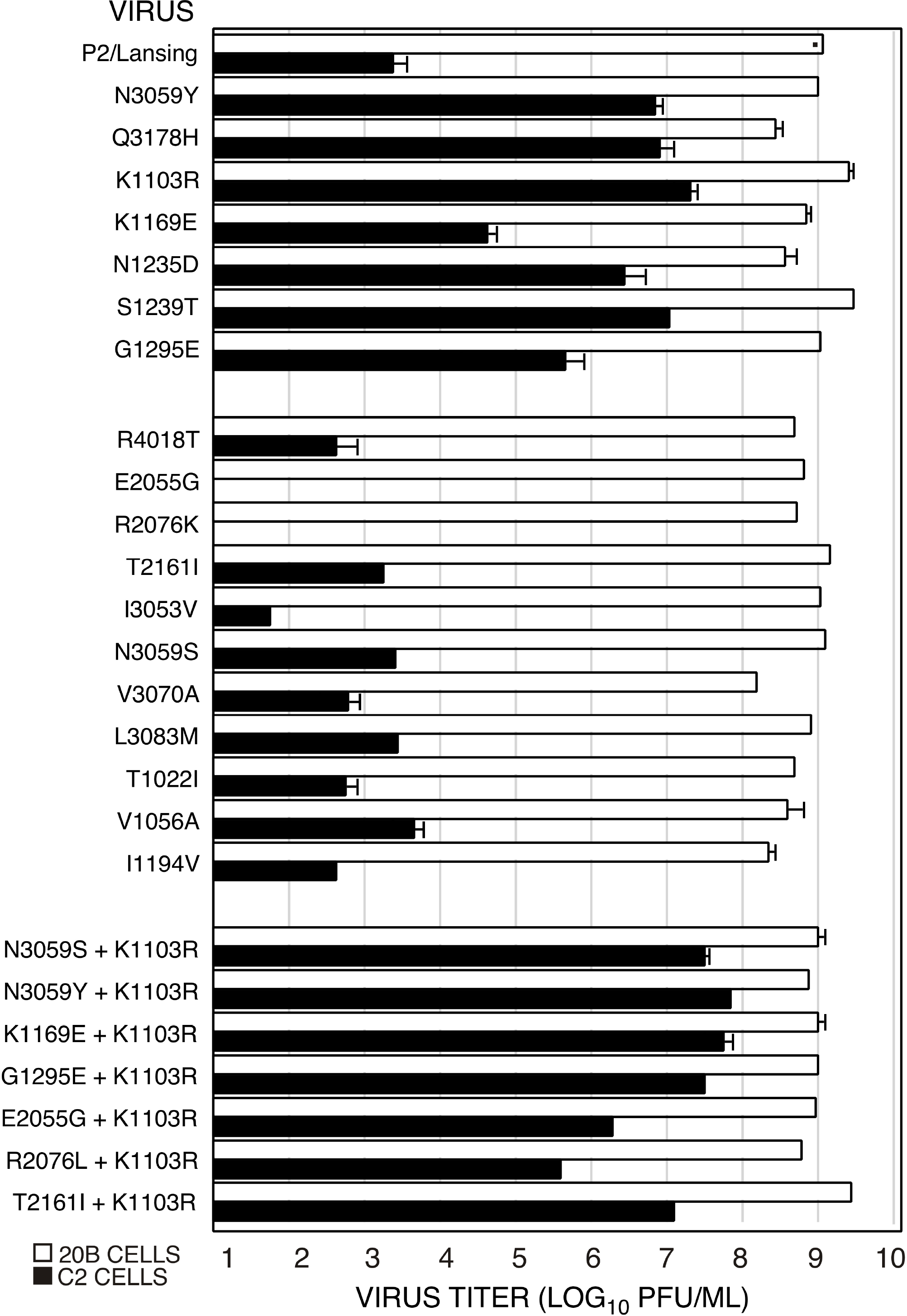
Growth of variants of P2/Lansing with single or double site-directed mutations on cells producing Pvr or nectin-2^Pvr (c’c”d)^. Virus titer was determined by plaque assays on monolayers of ⃞ 20B or ■C2 cells. The values are the averages of two or more independent viral constructs. For mutant viruses E2055G and R2076K, no plaques were observed on C2 cells at 10^-1^ fold dilution.

Multiple changes that were introduced individually into P2/Lansing did not confer the ability to replicate in C2 cells (Figure 4). Many of these changes were accompanied by K1103R in the original virus (Table 1). Additional viruses were therefore constructed that contain K1103R and one other amino acid change in the capsid. However, in no case was the titer in C2 cells improved by the presence of two amino acid changes in the capsid (Figure 4). In two viruses with two changes, E2055G + K1103R and R2076L + K1103R, viral titers on C2 cells were lower than in viruses containing only K1103R. Because viruses containing the single changes E2055G or R2076K did not form plaques on C2 cells, we conclude that other mutations must be necessary to permit growth of the parent viruses containing these substitutions. Unfortunately, it was not practical to test all possible combinations of amino acid changes.

### Adapting mutations improve binding of P2/Lansing to chimeric Pvr-nectin-2

Binding assays were done to determine whether mutations in P2/Lansing that enable it to replicate in C2 cells allow the virus to bind to the nectin-2^Pvr(c'c"d)^ receptor. In contrast to P2/Lansing, which cannot bind to C2 cells, viruses containing capsid substitutions K1103R, G1295E, S1239T, N3059Y, Q3178H, and K1169E can bind to C2 cells in a cell concentration-dependent manner (Figure 5). The extent of binding of these mutant viruses to C2 cells was comparable to that observed with 20B cells. These results show that certain amino acid changes in the viral capsid allow P2/Lansing to attach to the nectin-2^Pvr(c'c"d)^ receptor, permitting virus entry and replication.

**Figure 5:**
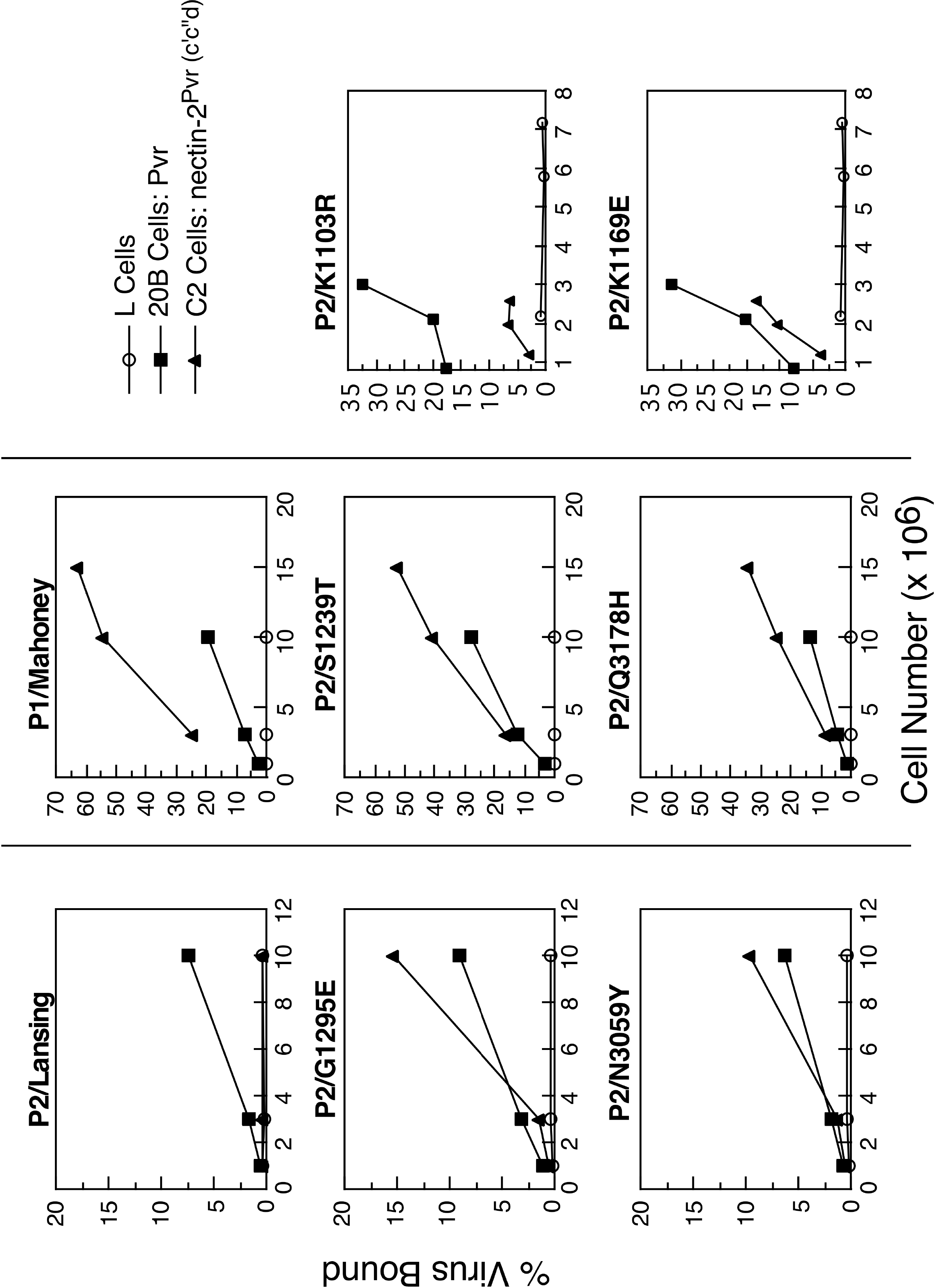
Binding assay of wild-type and mutant poliovirus on cells producing Pvr or nectin-2^Pvr(c’c”d)^. Radiolabeled poliovirus was incubated with ⃝ L cells, ■20B cells, and ▲ C2 cells at 37°C for 1 hour. Specific binding is expressed as a percentage of the total input virus.

### The VP1 BC loop is required for viral replication in C2 cells

The results of previous studies have demonstrated that the BC loop of VP1 can regulate receptor recognition and host range (Racaniello, 2013). Here we find that an amino change at the carboxy terminus of the VP1 BC loop, K1103R, can allow P2/Lansing to bind the nectin-2^Pvr(c'c"d)^ and was identified in 21 out of 24 virus isolates. To assess the importance of the VP1 BC loop in viral growth on C2 cells, we studied viral replication of variants of P1/Mahoney and P2/Lansing with changes in the VP1 BC loop.

The results demonstrate that P1/Mahoney requires an intact VP1 BC loop to replicate in C2 cells: a variant of P1/Mahoney from which the VP1 BC loop was deleted, PV-Δ9 virus, did not form plaques on these cells (Figure 6). In contrast, PV-414 virus, in which the VP1 BC loop of P1/Mahoney is substituted with the corresponding sequence from P2/Lansing, replicates to a high titer on C2 cells. The titer of PV-414 virus on C2 cells was greater than the titer of P1/Mahoney, suggesting that VP1 BC loop of P2/Lansing is more effective in enabling growth on C2 cells. These results suggest that P1/Mahoney requires the VP1 BC loop for interaction with nectin-2^Pvr(c'c"d)^.

**Figure 6:**
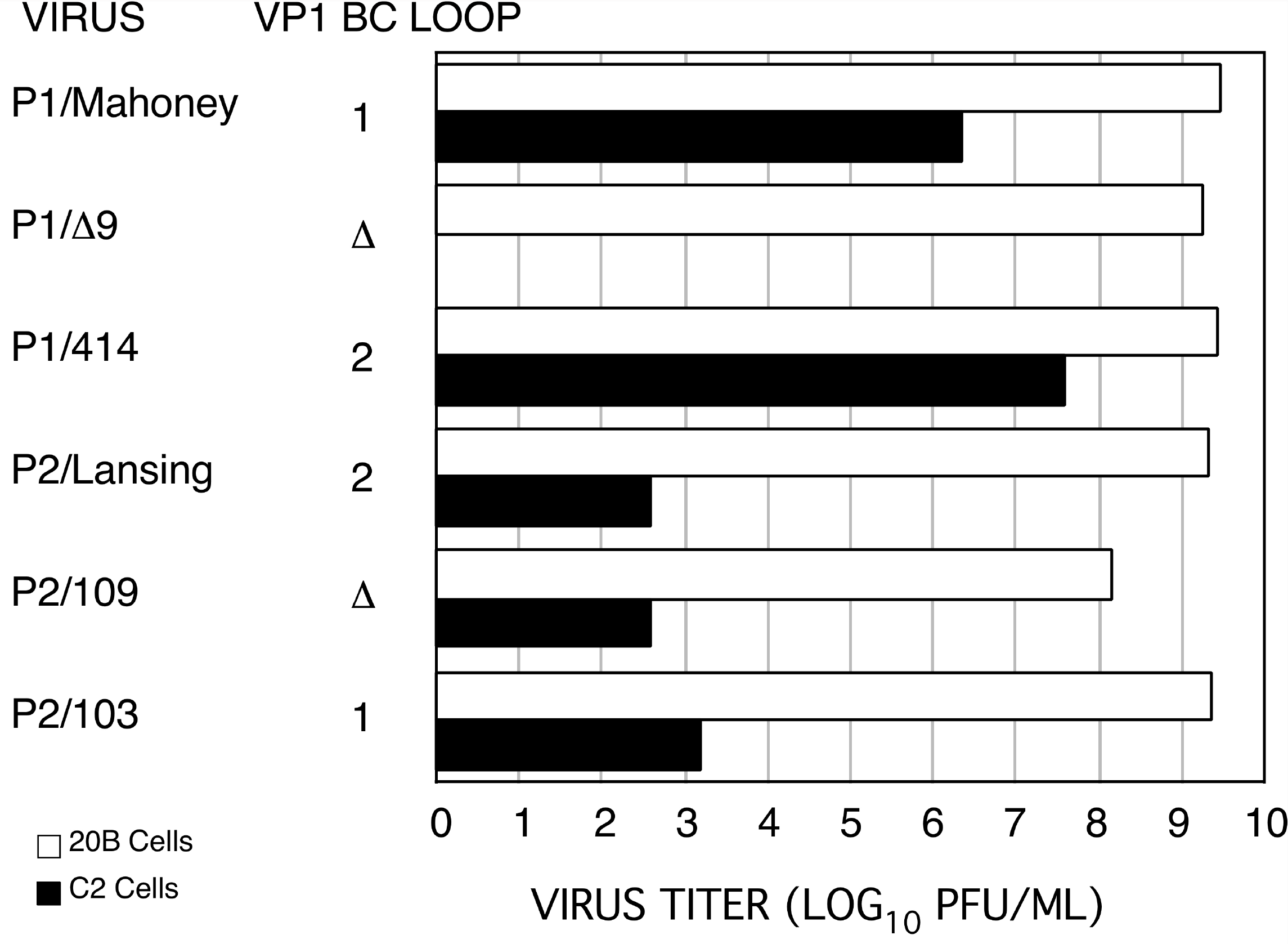
Yields of VP1 BC loop variants of polioviruses P1/Mahoney and P2/Lansing on cells producing Pvr or nectin-2^Pvr (c’c”d)^. Virus titer was determined by plaque assays on monolayers of ⃞20B or ⬛C2 cells. For P1/Δ9 virus, no plaques were observed on C2 cells at 10^-1^ fold dilution.

P2/Lansing cannot replicate in C2 cells, and deletion of the VP1 BC loop (P2/109 virus), or substitution with the corresponding sequence from P1/Mahoney (P2/103 virus), did not change this phenotype (Figure 6). Although the ability of P1/Mahoney to replicate in C2 cells depends on the type 1 VP1 BC loop. However, transfer of the P1/Mahoney VP1 BC loop to P2/Lansing is not sufficient to allow replication on C2 cells.

## Discussion

### Nectin-2^Pvr(c’c”d)^ is a functional receptor for poliovirus

Amino acid changes in the C’C”D region of Pvr domain 1 disrupt poliovirus binding (Aoki et al., 1994; Bernhardt et al., 1994b; Morrison et al., 1994), and models of the poliovirus-receptor complex based on cryoelectron microscopy and crystallographic data show that residues within the C’C”D region make extensive contact with the viral capsid (Belnap et al., 2000; Bostina et al., 2007; Bubeck et al., 2005; He et al., 2000; He et al., 2003; Strauss et al., 2015; Zhang et al., 2008). To determine whether this region is sufficient for virus binding, we constructed a poliovirus binding site in a murine homolog of Pvr nectin-2 by substituting a sequence encoding 31 amino acids of Pvr (amino acids 69-100) into the corresponding region. This chimeric molecule is topologically different from Pvr, as it is not recognized by a monoclonal antibody directed against the poliovirus binding site in Pvr domain 1 (Figure 1, data not shown), yet it is a functional receptor for poliovirus P1/Mahoney, but not for type 2 or type 3 polioviruses. The C’C”D region of Pvr in the context of the nectin-2 molecule is sufficient for viral entry only by P1/Mahoney.

Domain 1 of Pvr contacts the poliovirus P1/Mahoney canyon at two points, along the eastern and western walls of the canyon (Figure 7). The binding site on the eastern wall of the canyon runs in a north-south direction along the C-terminal extension of VP1. Other contacts of the receptor at this site include the C terminus of VP3 and a knob-like insertion in the B beta strand of VP3. The Pvr residues involved in contact on the eastern canyon wall are located in the CC’, C”D, and EF loops of domain 1. The contact site on the western wall of the canyon runs in a north-south direction along the H beta strand and GH loop of VP1. Pvr domain 1 also contacts the GH loops of VP1 and VP3, and the EF loop of VP2. At this site the contacts with Pvr include the FG, C’C” and DE loops, and the C” and G beta strands of domain 1. The C’C” loop of the receptor interacts with the GH loop of capsid protein VP1 (amino acids 116-119) and the N-terminal end of the D-strand contacts resides in VP3 (amino acids 58-60) and the C-terminus of VP1 (amino acids 293-397).

**Figure 7:**
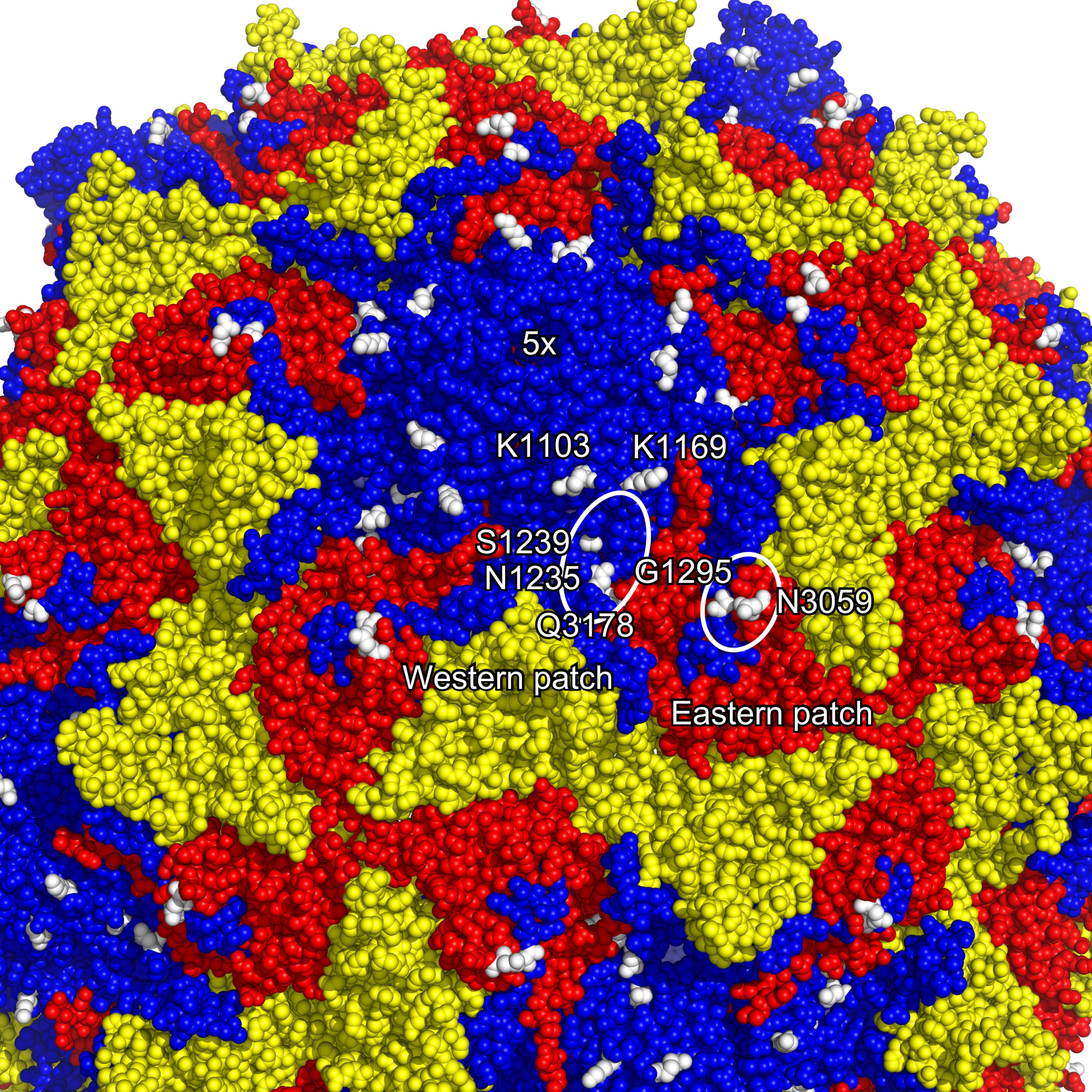
Location of amino acid changes in the viral capsid. Space-filling rendering of a poliovirus P2/Lansing pentamer viewed obliquely from the top. Color scheme: VP1 is blue, VP2 is yellow, and VP3 is red. VP4, which is internal, is not visualized. Single amino acid changes in VP1 and VP3 that allow poliovirus P2/Lansing to replicate on nectin-2^Pvr (c’c”d)^ producing cells are shown in white; residues in one structural unit are labeled. One five-fold axis of symmetry is labeled. Western and eastern contact sites on the canyon walls of Pvr on poliovirus P1/Mahoney are indicated by white ovals (Strauss et al., 2015). The ovals are oriented in the north-south direction on the capsid. The model is based on crystallographic data from Protein Data Bank file 1eah (Lentz et al., 1997).

Sites of Pvr outside of the C’C”D region that contact poliovirus are derived from nectin-2 in nectin-2^Pvr(c’c”d)^, yet, the recombinant receptor can bind to poliovirus P1/Mahoney. Viral capsid residues that interact with these receptor sites may be more tolerant of amino acid changes. Alternatively, the exact amino acid composition at these sites may not be as critical as that of the C’C”D region. Tertiary structures at these sites may be the major determinants of receptor recognition by the capsid. Although the overall amino acid identify between domains 1 of Pvr and nectin-2 is approximately 50%, the distribution of identical amino acid is not uniform throughout the domain; sites outside the C’C”D region account for much of the homology. Homolog-scanning mutagenesis of residues in these sites have been shown not to alter poliovirus P1/Mahoney binding and replication (Morrison et al., 1994). The observation that nectin-2^Pvr(c’c”d)^ cannot mediate cell entry of poliovirus serotypes 2 and 3 indicate that nectin-2 amino acids outside of the C’C”D region are not compatible with binding of these viruses.

### Capsid sequences that regulate receptor binding and entry

The inability of poliovirus P2/Lansing to replicate in cells that synthesize nectin-2^Pvr(c’c”d)^ and the subsequent identification of suppressor mutations affords an opportunity to study capsid residues involved in receptor interaction. The amino acid changes identified in this work are located along the canyon surface, at the protomer interface, and along the canyon perimeter (Figure 7). The results correlate with previous mutational analyses and structural models derived from cryoelectron microscopy and x-ray crystallographic data (Belnap et al., 2000; Bostina et al., 2007; Bubeck et al., 2005; He et al., 2000; He et al., 2003; Strauss et al., 2015; Zhang et al., 2008).

It has not been possible to select variants of P2/Lansing that can infect cells producing wild-type nectin-2 (unpublished results). It might be possible to isolate such variants beginning with P2/Lansing mutants adapted to replicate on C2 cells.

### Amino acid changes in the Pvr contact region on poliovirus

Alterations at capsid residues K1169, N1235, S1239, G1295, and N3059 permit P2/Lansing to bind the chimeric receptor (Figure 7). Changes at these residues may compensate for differences in the receptor chimera that prevent virus binding. Residue K1169 is located in the VP1 EF loop. This loop is at an interface between two fivefold-related protomers and in P1/Mahoney is part of the site on the western canyon wall that makes contact with the G strand of Pvr (Figure 7) (Strauss et al., 2015). Mutations at adjacent residues (E1168G and W1170R) enable P1/Mahoney to utilize mutant Pvr (Liao and Racaniello, 1997). In poliovirus P1/Mahoney, residue N1235 is located in the VP1 GH loop on the western canyon wall and contacts the Pvr C’C” loop, and residue S1239 is in the H strand and contacts the Pvr FG loop (Strauss et al., 2015). Nearby residues (A1231, L1234, and D1236) have been previously identified by *srr* (soluble receptor resistant) mutations as possible receptor contact sites (Colston and Racaniello, 1994). In P1/Mahoney, A1231 and L1234 contact the C’C” loop of Pvr (Strauss et al., 2015).

Residue G1295 is at the VP1 C-terminus on the eastern canyon wall, and contacts the Pvr CC’ loop (Figure 7) (Strauss et al., 2015). Amino acid N3059 is nearby and is part of the β-strand knob insert that contacts the Pvr EF loop (Strauss et al., 2015). This region is important in host range determination: mutations at the neighboring amino acid 3060 confer mouse neurovirulence to poliovirus P1/Mahoney (Couderc et al., 1994). Amino acid changes at positions 1295 and 3059 result in the lengthening of the side chains, an addition of a negative charge with G1295E, and the preservation of polarity with N3059Y. These alterations may enable poliovirus P2/Lansing to interact with the chimeric receptor effectively by possibly bridging a gap between the receptor-capsid interface.

None of the amino acids at positions 1169, 1235, 1239, 1295 and 3059 are identical in P2/Lansing selected for growth on C2 cells and in P1/Mahoney, which can infect C2 cells. In P3/Leon, which cannot replicate in C2 cells, nevertheless the amino acids at positions 1235 and 1295 are the same as in P2/Lansing selected for growth on C2 cells: D1235 and E1295.

### Mutations that influence poliovirus capsid dynamics

Receptor recognition may be influenced not only by residues in the virus-receptor interface, but also by amino acids that regulate capsid flexibility. In solution, the poliovirus capsid is structurally dynamic (Li et al., 1994; Roivainen et al., 1993), and virus binding to the receptor is inhibited at low temperatures by compounds that make the capsid structurally rigid (Dove and Racaniello, 2000). For example, amino acid change Q3178H, which broadens receptor utilization of P2/Lansing, maps to the same amino acid position as a *srr* alteration on P1/Mahoney and a *ts* suppressor in P3/Sabin (Colston and Racaniello, 1994; Filman et al., 1989), but does not appear to contact Pvr (Figure 7) (Strauss et al., 2015). *Srr* viruses with the Q3178L change less readily undergo conversion to noninfectious altered particles upon incubation with receptor. This substitution is thought to render the capsid more rigid, thus increasing its resistance to neutralization by soluble receptors. Conversely, the Q3178H change may cause greater capsid flexibility, thereby facilitating interactions with the less optimal nectin-2^Pvr(c’c”d)^ receptor.

In both P1/Mahoney, which can replicate in C2 cells, and in P3/Leon, which cannot replicate in these cells, the amino acids at position 3178 is glutamine, which was changed to histidine in P2/Lansing selected for growth on C2 cells.

Although amino acid S1239C contacts the Pvr FG loop in P1/Mahoney, it may also influence capsid flexibility by altering the structure of the hydrocarbon-binding pockets. In P1/Mahoney, a change at position 1241 is associated with the *srr* phenotype (Colston and Racaniello, 1994). The change (A1241V) may lead to a conformational shift in the hydrocarbon-binding pocket, resulting in increased resistance to receptor neutralization. In P2/Lansing, residue L1240 occupies the analogous position in lining the hydrophobic pocket (Lentz et al., 1997). By exerting its effects on the neighboring residue, S1239T may indirectly alter the interaction within the hydrocarbon-binding pocket. The changes may enable the virus to achieve the dynamic flexibility necessary for binding the chimeric receptor.

### K1103R and VP1 BC Loop

The change K1103R is present in 21 of the 28 mutants of P2/Lansing that were studied, suggesting that its presence is highly favored. This amino acid is located along the side of the canyon north wall, at the carboxyl terminal end of the VP1 BC loop (Figure 7) (Lentz et al., 1997). In poliovirus P1/Mahoney, residue K1103 does not make contact with the receptor, although the adjacent residue, D1102, contacts Pvr G strand. In poliovirus type 2 K1103 does not appear to contact Pvr (Zhang et al., 2008).

It is curious that both P1/Mahoney, which can replicate on C2 cells, and P3/Leon, which cannot replicate on these cells, have a lysine at position 1103. P2/Lansing also has a lysine at this position but it changes to arginine after selection for growth on C2 cells.

The three-dimensional structure of the VP1 BC loop differs significantly among the serotypes of poliovirus, and in poliovirus P2/Lansing it is structurally disordered (Lentz et al., 1997). The orientations of amino acid side-chains at residues K1103 and L1104 are reversed between polioviruses P1/Mahoney and P3/Sabin (Yeates et al., 1991). The structure of the carboxyl terminus of the P2/Lansing VP1 BC loop is more similar to that of P3/Sabin than P1/Mahoney (Lentz et al., 1997). The difference in the carboxyl terminus of VP1 BC loop may in part be due to the non-conserved amino acid sequence in the BC loop. Conversely, changes in the amino acid sequence that flank the BC loop are likely to result in a pronounced effect on the loop. Therefore K1103R may alter the conformation of neighboring structures, enabling P2/Lansing to recognize the chimeric receptor.

Another explanation for the mechanism of action of mutation K1103R is that it modulates capsid flexibility by changing the conformation of the VP1 BC loop. Realignment of the VP1 BC loop may alter the structure of neighboring DE and HI loops and create a new thermodynamic state at the five-fold axis of symmetry (Filman et al., 1989; Yeates et al., 1991). Amino acids at the amino and carboxyl termini of the VP1 BC loop may function as fulcra in transferring of energy from the relatively flexible BC loops to structures involved in receptor binding and viral uncoating. K1103R may make this process more efficient and compensate for the low affinity of P2/Lansing for the chimeric receptor. Consistent with this hypothesis, amino acid changes located at the amino terminal end of the VP1 BC loop (P1095S/T) suppress defects in Pvr at three different loci (Colston and Racaniello, 1995). This suppressor is not allele specific – it can suppress the effects of mutations at different positions - xand the amino acid change is postulated to affect the capsid globally by reducing the thermal requirement for virus binding to Pvr (Dove and Racaniello, 2000).

Our results show that the VP1 BC loop is required for binding of polioviruses P1/Mahoney and P2/Lansing to nectin-2^Pvr(c’c”d)^. This observation is surprising in light of the finding that the VP1 BC loop can be deleted without effecting the ability of either virus to bind Pvr and replicate in cultured cells (Martin et al., 1988; Murray et al., 1988). The VP1 BC loop is therefore not necessary for viral infection of cells producing high affinity, wild-type receptors, but is required for infection of cells synthesizing the low affinity chimeric receptor. Serotype 2 viruses poorly infect nectin-2^Pvr(c’c”d)^ cells even if a type 1 VP1 BC loop is present, indicating that the presence of this sequence is insufficient for complete complementation of the binding defects with nectin-2^Pvr(c’c”d)^. This observation further emphasizes the role of the carboxyl terminus (residue 1103) of the VP1 BC loop in chimeric receptor recognition.

Previous genetic analysis of the interaction of poliovirus with Pvr focused largely on serotype 1. In this report, we studied the binding of poliovirus type 2, and our findings extend our understanding of the poliovirus-receptor interaction, identify new sites that regulate receptor binding, and provide additional genetic confirmation for published structural models. We demonstrate that a chimeric protein made up of only a small portion of PVR—specifically, the C’C”D regions–in the context of a similar molecule permits viral infection. Serotypic differences in receptor utilization were revealed among the viruses and capsid changes were identified that enabled the virus to overcome the receptor defects. Analysis of these mutations corroborated the location of receptor binding sites predicted from previous genetic and structural data (Figure 7).

The identification of alterations in a receptor binding site on antigenic sites – the VP1 BC loop and the rim of canyon south wall (‘knob’) – raises interesting questions regarding the restricted antigenic variation observed among poliovirus in comparison to some other members of enterovirus family. Both amino acid changes identified on the ‘knob’ (G1295 and N3059) are to residues that occur in P3/Leon. Could participation in receptor recognition restrict amino acid variability at these sites needed to evade immune surveillance? (Rossmann, 1989). As P3/Leon cannot bind the chimeric receptor, what additional regions of the capsid are required for this virus? Additional genetic, biochemical, and structural analyses of the virus-receptor complex are needed to address these questions.

## Acknowledgements

We thank M. Girard and E. Wimmer for gifts of viruses PV-vΔ9 and PV-414 respectively, and Amy Rosenfeld for critical reading of the manuscript.

## References

Aoki, J., Koike, S., Ise, I., Sato-Yoshida, Y., Nomoto, A., 1994. Amino acid residues on human poliovirus receptor involved in interaction with poliovirus. J Biol Chem 269, 8431–8438.

Belnap, D.M., McDermott, B.M., Jr., Filman, D.J., Cheng, N., Trus, B.L., Zuccola, H.J., Racaniello, V.R., Hogle, J.M., Steven, A.C., 2000. Three-dimensional structure of poliovirus receptor bound to poliovirus. Proc Natl Acad Sci U S A 97, 73–78.

Bernhardt, G., Bibb, J.A., Bradley, J., Wimmer, E., 1994a. Molecular characterization of the cellular receptor for poliovirus. Virol. 199, 105–113.

Bernhardt, G., Harber, J., Zibert, A., DeCrombrugghe, M., Wimmer, E., 1994b. The poliovirus receptor: Identification of domains and amino acid residues critical for virus binding. Virology 203, 344–356.

Bostina, M., Bubeck, D., Schwartz, C., Nicastro, D., Filman, D.J., Hogle, J.M., 2007. Single particle cryoelectron tomography characterization of the structure and structural variability of poliovirus-receptor-membrane complex at 30 A resolution. J Struct Biol 160, 200–210.

Bubeck, D., Filman, D.J., Hogle, J.M., 2005. Cryo-electron microscopy reconstruction of a poliovirus-receptor-membrane complex. Nat Struct Mol Biol 12, 615–618.

Colston, E., 1995. Genetic determinants of the poliovirus-receptor interaction, Microbiology. Columbia University College of Physicians & Surgeons, New York.

Colston, E., Racaniello, V.R., 1994. Soluble receptor-resistant poliovirus mutants identify surface and internal capsid residues that control interaction with the cell receptor. EMBO J. 13, 5855–5862.

Colston, E.M., Racaniello, V.R., 1995. Poliovirus variants selected on mutant receptor-expressing cells identify capsid residues that expand receptor recognition. J Virol 69, 4823–4829.

Couderc, T., Guédo, N., Calvez, V., Pelletier, I., Hogle, J., Colbère-Garapin, F., Blondel, B., 1994. Substitutions in the capsids of poliovirus mutants selected in human neuroblastoma cells confer on the Mahoney type 1 strain a neurovirulent phenotype in mice. J. Virol. 68, 8386–8391.

Dove, A.W., Racaniello, V.R., 2000. An antiviral compound that blocks structural transitions of poliovirus prevents receptor binding at low temperatures. J Virol 74, 3929–3931.

Filman, D.J., Syed, R., Chow, M., Macadam, A.J., Minor, P.D., Hogle, J.M., 1989. Structural factors that control conformational transitions and serotype specificity in type 3 poliovirus. EMBO J. 8, 1567–1579.

Harber, J., Bernhardt, G., Lu, H.-H., Sgro, J., Wimmer, E., 1995. Canyon rim residues, including antigenic determinants, modulate serotype-specific binding of polioviruses to mutants of the poliovirus receptor. Virol. 214, 559–570.

Harrison, O.J., Vendome, J., Brasch, J., Jin, X., Hong, S., Katsamba, P.S., Ahlsen, G., Troyanovsky, R.B., Troyanovsky, S.M., Honig, B., Shapiro, L., 2012. Nectin ectodomain structures reveal a canonical adhesive interface. Nat Struct Mol Biol 19, 906–915.

He, Y., Bowman, V.D., Mueller, S., Bator, C.M., Bella, J., Peng, X., Baker, T.S., Wimmer, E., Kuhn, R.J., Rossmann, M.G., 2000. Interaction of the poliovirus receptor with poliovirus. Proc Natl Acad Sci U S A 97, 79–84.

He, Y., Mueller, S., Chipman, P.R., Bator, C.M., Peng, X., Bowman, V.D., Mukhopadhyay, S., Wimmer, E., Kuhn, R.J., Rossmann, M.G., 2003. Complexes of poliovirus serotypes with their common cellular receptor, CD155. J Virol 77, 4827–4835.

Hogle, J.M., Chow, M., Filman, D.J., 1985. Three-dimensional structure of poliovirus at 2.9 Å resolution. Science 229, 1358–1365.

Koike, S., Horie, H., Dise, I., Okitsu, H., Yoshida, M., Iizuka, N., Takeuthi, K., Takegami, T., Nomoto, A., 1990. The poliovirus receptor protein is produced both as membrane-bound and secreted forms. EMBO J. 9, 3217–3224.

Koike, S., Ise, I., Nomoto, A., 1991. Functional domains of the poliovirus receptor. Proc. Natl. Acad. Sci. USA 88, 4104–4108.

La Monica, N., Almond, J.W., Racaniello, V.R., 1987. A mouse model for poliovirus neurovirulence identifies mutations that attenuate the virus for humans. J. Virol. 61, 2917–2920.

Lentz, K.N., Smith, A.D., Geisler, S.C., Cox, S., Buontempo, P., Skelton, A., DeMartino, J., Rozhon, E., Schwartz, J., Girijavallabhan, V., O’Connell, J., Arnold, E., 1997. Structure of poliovirus type 2 Lansing complexed with antiviral agent SCH48973: comparison of the structural and biological properties of three poliovirus serotypes. Structure 5, 961–978.

Li, Q., Yafal, A.G., Lee, Y.M., Hogle, J., Chow, M., 1994. Poliovirus neutralization by antibodies to internal epitopes of VP4 and VP1 results from reversible exposure of these sequences at physiological temperature. J Virol 68, 3965–3970.

Liao, S., Racaniello, V., 1997. Allele-specific adaptation of poliovirus VP1 B-C loop variants to mutant cell receptors. J Virol 71, 9770–9777.

Martin, A., Wychowski, C., Couderc, T., Crainic, R., Hogle, J., Girard, M., 1988. Engineering a poliovirus type 2 antigenic site on a type 1 capsid results in a chimaeric virus which is neurovirulent for mice. EMBO J. 7, 2839–2847.

McCutchan, Pagano, J., 1968. J. Natl. Can. Inst. 41, 351–357.

Mendelsohn, C., Wimmer, E., Racaniello, V.R., 1989. Cellular receptor for poliovirus: Molecular cloning, nucleotide sequence and expression of a new member of the immunoglobulin superfamily. Cell 56, 855–865.

Morrison, M.E., Racaniello, V.R., 1992. Molecular cloning and expression of a murine homolog of the human poliovirus receptor gene. J. Virol. 66, 2807–2813.

Morrison, M.E., Yuan-Jing, H., Wien, M.W., Hogle, J.W., Racaniello, V.R., 1994. Homolog scanning mutagenesis reveals poliovirus receptor residues important for virus binding and replication. J. Virol. 68, 2578–2588.

Moss, E.G., O’Neill, R.E., Racaniello, V.R., 1989. Mapping of attenuating sequences of an avirulent poliovirus type 2 strain. J. Virol. 63, 1884–1890.

Moss, E.G., Racaniello, V.R., 1991. Host range determinants located on the interior of the poliovirus capsid. EMBO J. 5, 1067–1074.

Murray, M.G., Bradley, J., Yang, X.F., Wimmer, E., Moss, E.G., Racaniello, V.R., 1988. Poliovirus host range is determined a short amino acid sequence in neutralization antigenic site I. Science 241, 213–215.

Page, G.S., Mosser, A.G., Hogle, J.M., Filman, D.J., Rueckert, R.R., Chow, M., 1988. Three dimensional structure of the poliovirus serotype 1 neutralizing determinants. J. Virol. 62, 1781–1794.

Racaniello, V.R., 2013. Picornaviridae: The Viruses and Their Replication, in: Knipe, D.M., Howley, P.M. (Eds.), Fields Virology, 6th ed. Wolters Kluwer Health, Lippincott Williams & Wilkins, Philadelphia.

Racaniello, V.R., Baltimore, D., 1981. Cloned poliovirus complementary DNA is infectious in mammalian cells. Science 214, 916–919.

Rice, P., Longden, I., Bleasby, A., 2000. EMBOSS: the European Molecular Biology Open Software Suite. Trends Genet 16, 276–277.

Roivainen, M., Piirainen, L., Rysa, T., Narvanen, A., Hovi, T., 1993. An immunodominant N-terminal region of VP1 protein of poliovirion that is buried in crystal structure can be exposed in solution. Virology 195, 762–765.

Rossmann, M.G., 1989. The canyon hypothesis. Hiding the host cell receptor attachment site on a viral surface from immune surveillance. J Biol Chem 264, 14587–14590.

Rueckert, R.R., 1996. Picornaviridae: The viruses and their replication, in: Fields, B.N. (Ed.), Fields Virology. Lippincott-Raven, Philadelphia, pp. 609–654.

Selinka, H.-C., Zibert, A., Wimmer, E., 1991. Poliovirus can enter and infect mammalian cells by way of an intercellular adhesion molecule 1 pathway. Proc. Natl. Acad. Sci. USA 88, 3598–3602.

Shepley, M.P., Sherry, B., Weiner, H.L., 1988. Monoclonal antibody identification of a 100-kDa membrane protein in HeLa cells and human spinal cord involved in poliovirus attachment. Proc. Natl. Acad. Sci. USA 85, 7743–7747.

Strauss, M., Filman, D.J., Belnap, D.M., Cheng, N., Noel, R.T., Hogle, J.M., 2015. Nectin-like interactions between poliovirus and its receptor trigger conformational changes associated with cell entry. J Virol 89, 4143–4157.

Wigler, M., Pellicer, A., Silverstein, S., Axel, R., 1978. Biochemical transfer of single-copy eucaryotic genes using total cellular DNA as donor. Cell 14, 725–731.

Yeates, T.O., Jacobson, D.H., Martin, A., Wychowski, C., Girard, M., Filman, D.J., Hogle, J.M., 1991. Three-dimensional structure of a mouse-adapted type2/type 1 poliovirus chimera. EMBO J. 10, 2331–2341.

Zhang, P., Mueller, S., Morais, M.C., Bator, C.M., Bowman, V.D., Hafenstein, S., Wimmer, E., Rossmann, M.G., 2008. Crystal structure of CD155 and electron microscopic studies of its complexes with polioviruses. Proc Natl Acad Sci U S A 105, 18284–18289.

